# Maternal dietary deficiency in choline reduced levels of MMP-2 levels in blood and brain tissue of male offspring mice

**DOI:** 10.1101/2024.07.15.603575

**Authors:** Mitra Esfandiarei, Shawn G.U. Strash, Amanda Covaleski, Sharadyn Ille, Wei-dang Li, Nafisa M. Jadavji

**Affiliations:** Department of Biomedical Sciences, Midwestern University, Glendale, AZ, USA; Anesthesiology, Pharmacology, & Therapeutics, Faculty of Medicine, University of British Columbia, Vancouver, BC, Canada; Basic Medical Sciences, College of Medicine Phoenix, University of Arizona, Phoenix, AZ, USA; College of Osteopathic Medicine, Midwestern University, Glendale, AZ, USA; College of Pharmacy, Midwestern University, Glendale, AZ, USA; College of Dental Medicine, Midwestern University, Glendale, AZ, USA; College of Veterinary Medicine, Midwestern University, Glendale, AZ, USA; Department of Biomedical Sciences, Southern Illinois University, Carbondale, IL, USA; Department of Child Health, College of Medicine Phoenix, University of Arizona, Phoenix, AZ, USA; Department of Neuroscience, Carleton University, Ottawa, ON, Canada

**Keywords:** maternal nutrition, folic acid, choline, ischemic stroke, offspring, one-carbon metabolism

## Abstract

Ischemic stroke is one of the leading causes of disability and death globally, with a rising incidence in younger age groups. It’s well known that maternal diet during pregnancy and lactation is vital for the early neurodevelopment of offspring. One-carbon (1C) metabolism, including folic acid and choline, plays a vital role in closure of the neural tube *in utero*. However, the impact of maternal dietary deficiencies in 1C on offspring neurological function following ischemic stroke later in life remains undefined. The aim of this study was to investigate inflammation in blood and brain tissue of offspring from mothers deficient in dietary folic acid or choline. Female mice were maintained on either a control or deficient diets prior to and during pregnancy and lactation. When offspring were 3-months of age, ischemic stroke was induced. One and half months later blood and brain tissue were collected. We measured levels of matrix-metalloproteases (MMP)-2 and 9 in both plasma and brain tissue, and report reduced levels of MMP-2 in both, with no changes observed in MMP-9. This observation supports our working hypothesis that maternal dietary deficiencies in folic acid or choline during early neurodevelopment impact the levels of inflammation in offspring after ischemic stroke.

## 1. Introduction

Ischemic stroke is the leading cause of long-term neurological impairments worldwide, leading to disability and death [1–3]. An ischemic stroke occurs when there is a blockage in a blood vessel in the brain, an example is a thrombus. The factors that increase risk for ischemic stroke, including hypertension, obesity, and diabetes, are on the rise [4–6]. Nutrition is linked to the onset of these aforementioned risk factors [7] and is a modifiable risk factor for ischemic stroke [8,9]. Previous literature has established a link between one-carbon (1C) metabolism deficiency and worse post-ischemic stroke outcome [10–17]. Folic acid and choline are the main components of 1C and are well known for their role in the closure of the neural tube of the developing fetus during pregnancy[18].

During pregnancy there is increased demand for maternal folic acid and choline, which is often obtained through the diet [19,20]. Both folic acid and choline play an important role in nucleotide synthesis, DNA repair, and methylation through their involvement in 1C [21]. These cellular processes are pivotal during pregnancy and important during normal neurodevelopment. The role of maternal folic acid and choline is well established for *in utero*, but less understood for offspring neurological function after birth.

The Developmental Origins of Health and Disease (DOHaD) theory suggests that prospective chronic diseases are programmed *in utero* [22,23]. In offspring, maternal dietary deficiencies have been associated with modified neural tube closure [24,25] and neurocognitive development [26–29]. Epidemiological studies have demonstrated the effect of maternal diet on lifelong cardiovascular and neurological function [30]. Female and male offspring from moms deficient in folic acid or choline have worse outcome after ischemic stroke including impaired behavior in both 4.5 and 11.5 month old groups [14,15]. We also report reduced blood flow in female offspring [16]. In these studies when investigating mechanisms, we reported reduced levels of apoptosis, response to hypoxia, neuroinflammation, when measured 1.5 months after ischemic damage was induced [14,15]. In this study, we wanted to investigate other mechanisms, such as inflammation at the same timpoint.

Matrix-metalloproteases (MMP) have been identified as proteins that become concentrated in areas of tissue undergoing response to injury and acute inflammation [31,32]. MMP’s have been correlated with a variety of diseases related to chronic inflammation or neurological disorders, such as ischemic stroke [31]. Following injury, increased expression of MMPs has been related to increased blood brain barrier (BBB) permeability due to disruption of the tight junctions and adherents’ junctions in the BBB. This increased permeability results in further exacerbation of potential CNS damage [9,33]. It is reported that mice with reduced production of MMP-9 (type of gelatinase MMP, along with MMP-2) following stroke presented with less cerebral injury compared to control mice [34]. A separate study also found that mice with increased production of MMP-9 had developed exacerbated cerebral injury compared to control groups [35]. These findings suggest the possibility of MMP’s being a potential therapeutic target for ischemic injuries.

Maternal dietary deficiencies in folic acid or choline impact offspring stroke outcome in female and male offspring [14–16], however the mechanism through which this occurs it not well understood. The aim of this study was to measure the levels of inflammatory markers MMP-2 and -9 post-stroke in the brain and plasma of female and male offspring from mothers deficient in dietary folic acid or choline during pregnancy and lactation.

## 2. Materials and Methods

All experiments were conducted in accordance with the guidelines of the National Institutes of Health Guide for the Care and Use of Laboratory Animals (8^th^ ed., 2011) and University Institutional Animal Care Users Committee (IACUC 2983) in February 2020. Female and male C57/BL6J (RRID: IMSR_JAX:000664) mice were obtained from Jackson Laboratories (Bar Harbor, ME, USA). All mice were housed in a controlled environment with a 12-hour light/dark cycle, a temperature of 22–23°C, a relative humidity of 55% to 60%, and free access to food and water. Male and female offspring were generated from breeding pairs.

Experimental manipulations are summarized in **Figure 1**. Two-month-old female mice were habituated for 7 days before they were fed on either control (CD, Envigo, Indianapolis, IN, Catalog: # TD.190790), folic acid (FADD, Envigo, Indianapolis, IN, Catalog: # TD.01546), or choline deficient diets (ChDD, Envigo, Indianapolis, IN, Catalog: # TD.06119). All diets contained 1% succinylsulfathia-zole to prevent folate synthesis by intestinal flora. The CD represented adequate levels of both folic acid and choline that would be found in a conventional diet, which were determined from previous literature and experimentation [36–39]. The deficient diets consisted of lower levels of folic acid or choline compared to the control diet. The CD contained 2 mg/kg of folic acid and 1,150 mg/kg of choline bitrate, whereas the FADD contained only 0.3 mg/kg and the ChDD only had 300 mg/kg of choline bitrate. The dams were maintained on the diets 4 weeks prior to pregnancy, during pregnancy, and lactation. Once female and male offspring were weaned from their mothers, they were maintained on a CD for the duration of the experiment. At 2 months of age, the offspring were subjected to ischemic stroke, using the photothrombosis model to the sensorimotor cortex. Four and half weeks after damage blood and brain tissue were collected for further analysis.

**Figure 1.**
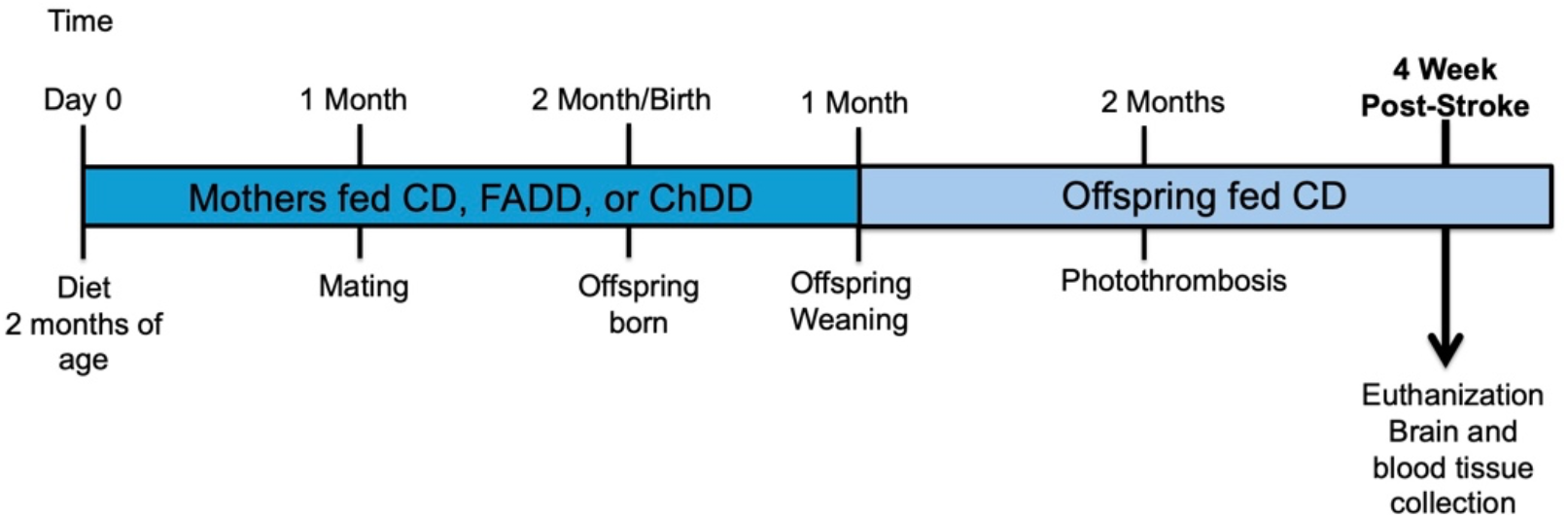
Experimental timeline of study. At 2-month-of age female mice were placed on control (CD), folic acid (FADD), or choline deficient (ChDD) diets. Females were maintained on diets for duration of pregnancy and lactation until the offspring were weaned. Once female and male offspring were weaned, they were maintained on the CD. At 2 months of age, the offspring were subjected to ischemic stroke via a unilateral photothrombosis model. At 3.5 months, female and male offspring were euthanized, and blood and brain tissues were collected for analysis.

### 2.1 Photothrombosis

At 2 months of age, all female and male offspring were subjected to photothrombosis to induce a unilateral ischemic stroke in the sensorimotor cortex. They were anesthetized with isoflurane (Henry Schein, Melville, NY, USA, Catalog: #NDC-11695-6776-1,) (1.5%) in a 70:30 nitrous oxide: oxygen mixture. Core body temperature was monitored with a rectal thermometer (Harvard Apparatus, Holliston, MA) and maintained at 37 ± 0.2ºC using a heating blanket. The photosensitive Rose Bengal dye (10mg/kg; (Millipore Sigma, Burlington, MA, USA, Catalog: #198250-25G) was injected intraperitoneally 5 minutes prior to irradiation. A 532 nm green laser Beta Electronics, Irvine, CA, USA (#MGM20 (20-25mW)) was placed 3 cm above the animal and directed to the sensorimotor cortex (mediolateral + 0.24mm) [10,12–15,40,41] for 15 min. All animals received 0.1 mg/kg of Burprenorphine HCl after damage to assist with post-operative pain. After the completion of surgery animals were placed in home singly housed. Animals were provided mash and hydrogel for up to one-week post-stroke and house individually.

### 2.2 Brain tissue collection, processing, and sectioning

At 4.5 weeks after stroke, female and male offspring mice were anesthetized with an overdose of CO_2_. Brain tissue was dissected from skull and placed into 4% paraformaldehyde overnight. Solution was replaced to 10% sucrose and then 20% sucrose, each change of solution lasted overnight. Once brain tissue was ready to be sectioned, it was frozen and mounted. Then frozen brain tissue was sectioned using a Thermo HM550 cryostat (Fisher Scientific, Hampton, NH, USA) at a thickness of 30μm and sections were slide mounted on microscope slides in serial order. Microscope slides were stored at -80°C until analysis.

### 2.3 Immunofluorescence

Brain tissue was used for immunofluorescence analysis to assess inflammation, particularly levels of MMP-2 and 9. Primary antibodies used included, MMP-2 (1:100, AbCam, Cambridge, MA, USA, Catalog: #ab97779, RRID: AB_10696122) and MMP-9 (1:500, AbCam, Cambridge, MA, USA, Catalog: #ab228402, RRID: AB_2910609). All brain sections were stained with a marker for neuronal nuclei (NeuN, 1:200, AbCam, Cambridge, USA, Catalog: # ab104224, RRID: AB_10711040). Primary antibodies were diluted in 0.5% Triton X diluted in phosphate buffered saline and incubated with brain tissue overnight at 4°C. Three rinses were conducted between each step using phosphate buffered saline, each rinse was 5 minutes in length. The next day, brain sections were incubated with secondary antibodies Alexa Fluor 488 and 555 (Cell Signaling Technologies Danvers, MA, USA, Catalog: # 4408 and 4413) at room temperature for 2 hours. Brain tissue was also stained with 4’, 6-diamidino-2-phenylindole (DAPI) (1:10000, Thermo Fisher Scientific, Waltham, MA, Catalog: # EN62248). Tissue sections were cover slipped with Fluromount and stored at 4°C in the dark until visualization. The staining was visualized using an Revlove microscope (Echo, San Diego, CA) and all images were collected at the magnification of 200X.

In brain tissue within the ischemic core region, co-localization of MMP-2 or MMP-9 with NeuN labelled neurons were counted and averaged per animal. Images were merged, and a positive cell was indicated by colocalization of the antibodies of interest located within a defined cell. Cells were distinguished from debris by identifying a clear cell shape and intact nuclei (indicated by DAPI and NeuN) under the microscope. All cell counts were conducted by three individuals blinded to treatment groups. Using ImageJ, the number of positive cells were counted in three brain sections per animal, a total of 3 to 4 animals were analyzed per group. For each section, three brain sections were analyzed per animal. Within each brain section three subsections were measured. The number of positive cells were averaged for each animal.

### 2.5 ELISA

Blood was collected in EDTA-coated tubes from experimental animals at time of euthanasia using cardiac puncture. Samples were spun down at 7000G for 7 minutes at 4°C and plasma was stored at -80°C until analysis. The mouse MMP-2 ELISA Kit (Abcam Inc, Cambridge, MA, USA, Catalog: #ab254516) and the MMP-9 ELISA Kit (Abcam Inc, Cambridge, MA, USA, Catalog: #ab253227) were used for the quantitative measurement of MMP-2 and MMP-9 in mouse plasma, respectively, according to the manufacturer’s instructions. In brief, the plasma samples were diluted according to the recommended dilution factor provided in the kit protocol, and 50 μL from each sample was added to each well. Then, 50 μL antibody cocktail was added to each well. A substrate solution 3,3’,5,5’-Tetramethylbensidine (TMB) was applied for visualization. The absorbance at 450nm was measured using a BIOTEK EPOCH (Agilen, Sanata Clara, CA) microplate reader.

### 2.6 Data Analysis and Statistics

GraphPad Prism 10.0 was used to analyze cell counts and ELISA data. In GraphPad Prism, D’Agostino-Pearson normality test was performed prior to two-way ANOVA analysis for all data. The two-way ANOVA compared the mean measurement of both sex and maternal dietary group for ELISA and immunofluorescence staining. One-way ANOVA analysis was used to compare maternal dietary differences when there were no offspring sex differences. Significant main effects of two- and one-way ANOVAs were followed up with Tukey’s post-hoc test to adjust for multiple comparisons. All data are presented as mean ± standard error of the mean (SEM). Statistical tests were performed using a significant *P* value of 0.05.

## 3. Results

### 3.1. Plasma levels of MMP-9 and MMP-2

Four and half weeks after ischemic stroke, we measured levels of MMP-2 and MMP-9 in plasma isolated from female and male offspring from mothers maintained on CD, ChDD, or FADD prior to and during pregnancy, and lactation. Offspring MMP-2 levels were affected by maternal diet (Figure 2A, F(_2,73_) = 5.82, p = 0.007). Male ChDD mice had lower levels of MMP-2 compared to FADD mice (p = 0.024). MMP-2 levels were not impacted by sex (F(_1,73_) = 2.20, p = 0.13) nor was there an interaction between sex and maternal diet (F(_2,73_) = 1.19, p = 0.031). The maternal diet offspring were exposed during *in utero* and lactation did not impact MMP-9 levels (Figure 2B; F(_2,73_) = 0.14, p = 0.867), sex (F(_1,73_) = 0.97, p = 0.97) or interaction between maternal diet and sex (F(_2,73_) = 0.82, p = 0.45).

**Figure 2.**
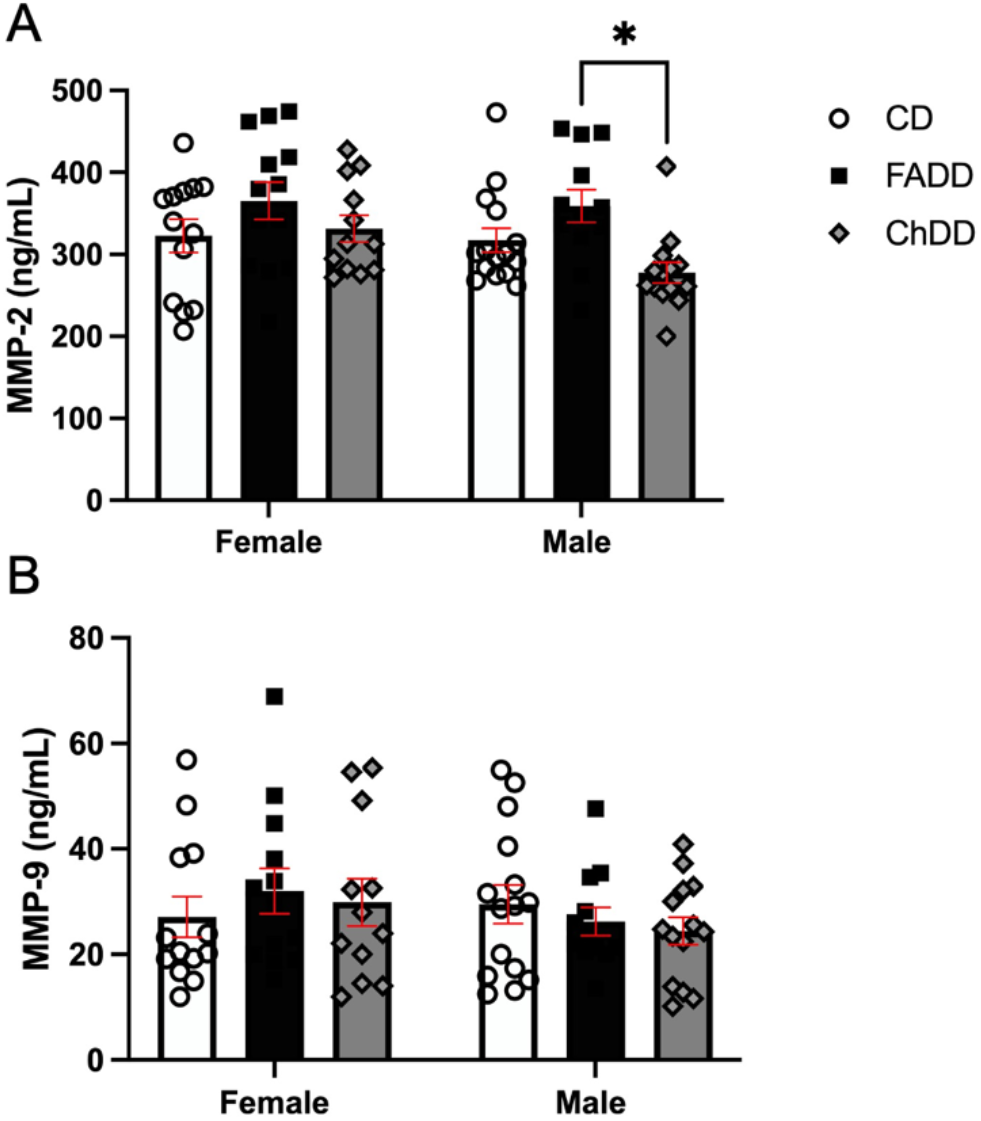
Impact of maternal diet deficient in folic acid (FADD) or choline (ChDD) and control (CD) during pregnancy and lactation on levels of matrix-metalloproteases (MMP) -2 (A) and 9 (B) in plasma of female and male offspring 1 month after ischemic stroke. Mean ± SEM of 10 to 14 animals per group. * p < 0.05, Tukey’s pairwise post-hoc analysis.

### 3.2. Brain Levels of MMP-2 and MMP-9

To assess inflammation in brain tissue we measured neuronal levels of MMP-2 and MMP-9 within the ischemic damage region. Representative images of MMP-2 and NeuN staining are shown in Figure 3A. The tissue was collected four and half weeks after ischemic stroke. Maternal diet did impact MMP-2 levels (Figure 3B, (F(_2, 14_) = 5.00, p = 0.02). There were no significant pairwise comparison differences. Furthermore, there was no impact of offspring sex (F(_1,14_) = 0.82, p = 0.95) or interaction between maternal diet and offspring sex (F(_2,14_) = 1.09, p = 0.37).

**Figure 3.**
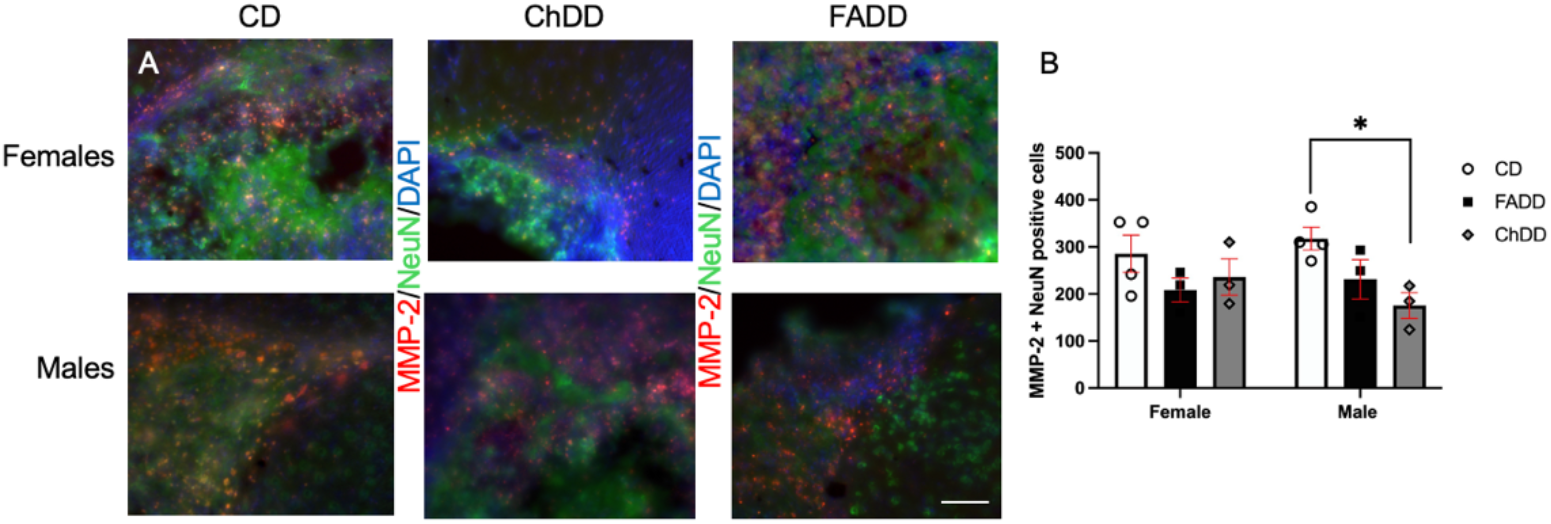
Impact of maternal diets deficient in folic acid (FADD) or choline (ChDD) and control diet (CD) during pregnancy on lactation on neuronal levels of matrix-metalloproteases (MMP-2) in ischemic brain tissue of female and male offspring one month after ischemic stroke. Representative images of MMP-2 immunofluorescence colocalization with neuronal nuclei (NeuN) and 4’,6-diamidino-2-phenylindole (DAPI) (A) and semi-quantitative quantification of MMP-2 levels within ischemic region of brain tissue. Mean ± SEM of 3 to 4 animals per group. * p < 0.05, Tukey’s pairwise post-hoc analysis (one-way ANOVA analysis between males and significant maternal diet main effect).

Since there was no sex difference between offspring, using one-way ANOVA we looked at the impact of maternal diet separately in female and male offspring. There was no impact of maternal diet on female levels of MMP-2 in brain tissue (F (_2, 7_) = 1.183, p = 0.361). Interestingly we see reduced levels of MMP-2 in male brain tissue because of maternal diet (F(_2, 7_) = 5.734, p = 0.034).). Specifically, between ChDD and CD offspring (p = 0.023; Figure 3B).

Representative images of MMP-9 and NeuN staining are shown in Figure 4A. MMP-9 levels were measured in brain tissue within the ischemic region, there was no impact of maternal diet (Figure 4B, F(_2, 11_)= 1.33, p = 0.30), offspring sex (F(_1, 11_)= 0.23, p = 0.64), or interaction between maternal diet and offspring sex (F(_2, 11_)= 0.71, p = 0.71).

**Figure 4.**
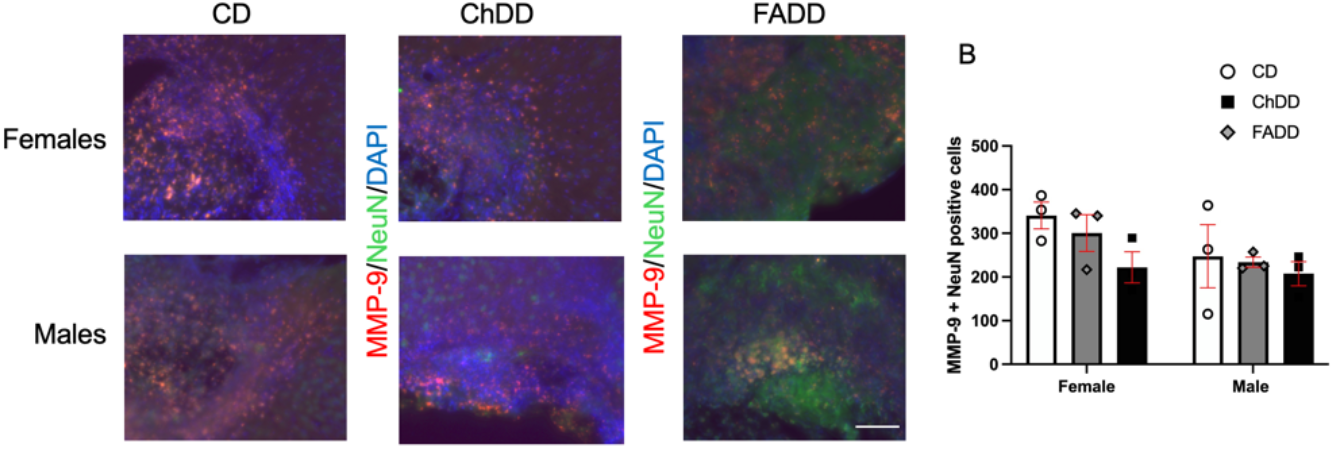
Impact of maternal diets deficient in folic acid (FADD) or choline (ChDD) and control diet (CD) during pregnancy and lactation on neuronal levels of MMP-9 in ischemic brain tissue of female and male offspring one month after ischemic stroke. Representative images of matrix-metalloproteases (MMP-9) immunofluorescence colocalization with neuronal nuclei (NeuN) and 4’,6-diamidino-2-phenylindole (DAPI) (A) and semi-quantitative quantification of MMP-9 levels within ischemic region of brain tissue. Mean ± SEM of 3 to 4 animals per group.

## 4. Discussion

Ischemic stroke leads to negative health outcomes around the world [1–3] Nutrition is a risk factor for ischemic stroke [8,9]. Previous literature has identified a link between post-stroke outcome and 1C metabolism [10–13,17,40]. The DOHaD theory suggests that prospective chronic diseases are programmed *in utero*-giving rise to programming of offspring cardiovascular, metabolic, and neuroendocrine dysfunction [22,23]. Despite impressive evidence of the importance of the maternal environment for fetal growth and development *in utero*, there is not much information available on the impact on offspring as they age. Nutrition is a modifiable risk factor [8] and can also impact outcome after ischemic stroke [12–15,42–44]. Low dietary levels of maternal folic acid or choline worsen stroke outcomes in female and male offspring [14–16]. This study aimed to investigate whether pro-inflammatory MMPs play a role in the impact of reduced maternal dietary levels of folic acid or choline on post-stroke outcomes in female and male offspring. We report that MMP-2 levels were decreased in the brain tissue and plasma of our choline deficient diet male offspring following ischemic stroke. Whereas MMP-9 levels were unaffected in treatment groups.

MMPs have been established as potential therapeutic targets after ischemic stroke due to their importance and critical role in the inflammation cascade. Decreased production of MMP following ischemic stroke has already been shown to result in decreased injury to the brain and blood brain barrier in mice [34,35]. Interestingly, our results show that male offspring from maternal mice with choline deficient diets have decreased MMP-2 levels present in their plasma and brain tissue following ischemic stroke. It is unexpected that the treatment groups who were exposed to maternal deficiencies of choline or folic acid during *in utero* and early postnatal development are seemingly able to correct for ischemic stroke more efficiently than their 1C sufficient control group comparison.

MMPs are inflammatory molecules leading to the disruption of barrier-like cellular structures such as tight junctions, adherents’ junctions, blood brain barrier dysfunction. Like other molecules in the inflammatory cascade, these molecules are important in biological processes such as immune support and wound healing, but when these molecules gather in excess, we see the negative consequences of inflammation. MMPs have been shown to be important in protective and beneficial inflammatory responses during normal physiological processes such as bone remodeling and angiogenesis [45]. Dietary deficiencies and their impact on MMP expression and activity in the brain have been studied extensively. Previous studies have shown that deficiencies in vitamin B (B6, B12, & folate) can elevate homocysteine levels, which in turn can increase MMP activity, contributing to neurovascular and brain damage [46,47]. Deficiencies in vitamin E and other antioxidants like selenium can also results in oxidative stress, increasing MMP activity in the brain [48,49].

In our previous reports, we have shown that offspring from moms deficient in folic acid or choline have shown worse outcome after ischemic stroke including impaired behavior in both 4.5 and 11.5 months of age [14,15]. We also reported reduced levels of apoptosis, response to hypoxia, neuroinflammation at 1.5 months after ischemic injury [14,15]. Hence, the observation of reduced MMP levels in offspring after ischemic stroke might have been expected. Since in the brain, MMP-2 drives neuronal apoptosis and breakdown of specific substrates such as dystroglycan, a transmembrane receptor involved in anchoring of astrocyte end feet to the basement membrane via laminin binding. Furthermore, cellular influx into areas of inflammation is regulated by MMP-2 activity [50]. The observed decrease in MMP-2 levels in ChDD mice can be due to the activation of compensatory pathways to counterbalance the deficiency, leading to an adaptive reduction in MMP expression. In addition, maternal protein deficiency could induce epigenetic modifications in the offspring, altering the expression of genes involved in inflammation and MMP regulation. Dietary deficiencies might also affect the development of the offspring’s immune system, resulting in a less robust inflammatory response and consequently lower MMP levels. Another possibility is that the offspring might develop protective adaptations against inflammation and oxidative stress due to chronic maternal deficiency during the developmental stage, leading to reduced MMP levels post-stroke.

From an experimental design perspective, these plasma and brain samples were taken four and half weeks after ischemic stroke was induced. Results could have potentially differed if samples were taken at different points in time, such as sample collection right after ischemic stroke. Potential future studies could include analyzing the same samples across different time frames and assessing if different levels of MMPs are noticed at different points in time. Previous research measured MMP levels at different times following cardiac ischemia. They noticed that MMP levels are increased in proportion to level of ischemia at the initial onset of ischemia (acute myocardial infarction vs stable coronary artery disease), but later decrease to below less severe ischemic levels [51]. This could potentially explain the results of our study. The diet deficient treatment groups may have undergone more severe ischemia than the control groups, having more MMPs initially but could potentially still have lower levels of MMP a short time later. The previously mentioned study regarding heart ischemia took their samples 3 months post ischemic injury, our results show similar findings at the 1.5-month mark.

## 5. Conclusions

We report reduced levels of MMP-2 in brain and plasma tissue of offspring one month after undergoing induced ischemic stroke from mothers deficient in choline. Previous research has shown that reduced levels of MMPs are associated with less severe injury and complications following cerebral ischemia. However, it would be an oversimplification to classify MMPs as inherently deleterious. MMPs are crucial players during tissue remodeling, repair, and healing processes, but their overexpression and uncontrolled activity can be detrimental, leading to excessive degradation of extracellular matrix components, contributing to inflammatory and pathological conditions. Further research is necessary to elucidate the precise benefits and adverse consequences of varying levels of MMPs in the brain of offspring from mothers with dietary deficiencies.

Additionally, studying MMP expression at different time points post-stroke is crucial to understanding their dynamic roles and impacts on neurological recovery and damage. Furthermore, MMP-2 is constitutively expressed in several tissues and is regulated by tumor necrosis factor-α under the influence of NF-κB transcription factor [50] – investigating these levels in offspring brain tissue is also a future goal. As well as inhibiting MMP-2 function may provide some clues to the role of this inflammatory marker on offspring stroke outcome.

## Supporting information

Supplementary Materials: Table S1: Mean and Standard Error of Mean data.

## Supplementary Materials

Table S1: Mean and Standard Error of Mean data.

## Author Contributions

Conceptualization, M.E. and N.M.J.; methodology, M.E., W.L., N.M.J.; validation, N.M.J., W.L.; formal analysis, S.G.U.S, A.C., S.I., N.M.J.; investigation, N.M.J.; resources, M.E., W.L., N.M.J.; data curation, N.M.J.; writing—original draft preparation, S.G.U.S., W.L., N.M.J; writing—review and editing, M.E., S.G.U.S., W.L., A.C., S.I., N.M.J; visualization, N.M.J.; supervision, M.E., N.M.J.; project administration, M.E., N.M.J.; funding acquisition, M.E., N.M.J. All authors have read and agreed to the published version of the manuscript.

## Institutional Review Board Statement

The animal study protocol was approved by the Institutional Review Board of MIDWESTERN UNIVERSITY (2983, February 24, 2020).

## Informed Consent Statement

Not applicable.

## Data Availability Statement

Not applicable.

## Acknowledgments

Not applicable.

## References

1. Prust, M.L.; Forman, R.; Ovbiagele, B. Addressing Disparities in the Global Epidemiology of Stroke. Nat Rev Neurol 2024, 20, 207–221, doi:10.1038/s41582-023-00921-z.

2. Campbell, B.C.V.; Khatri, P. Stroke. Lancet 2020, 396, 129–142, doi:10.1016/S0140-6736(20)31179-X.

3. Donkor, E.S. Stroke in the 21st Century: A Snapshot of the Burden, Epidemiology, and Quality of Life. Stroke Res Treat 2018, 2018, 3238165, doi:10.1155/2018/3238165.

4. Sarma, S.; Sockalingam, S.; Dash, S. Obesity as a Multisystem Disease: Trends in Obesity Rates and Obesity-Related Complications. Diabetes, Obesity and Metabolism 2021, 23, 3–16, doi:10.1111/dom.14290.

5. Mills, K.T.; Stefanescu, A.; He, J. The Global Epidemiology of Hypertension. Nat Rev Nephrol 2020, 16, 223–237, doi:10.1038/s41581-019-0244-2.

6. Boehme, A.K.; Esenwa, C.; Elkind, M.S.V. Stroke Risk Factors, Genetics, and Prevention. Circ Res 2017, 120, 472–495, doi:10.1161/CIRCRESAHA.116.308398.

7. Mozaffarian, D. Dietary and Policy Priorities for Cardiovascular Disease, Diabetes, and Obesity. Circulation 2016, 133, 187–225, doi:10.1161/CIRCULATIONAHA.115.018585.

8. Hankey, G.J. Nutrition and the Risk of Stroke. The Lancet Neurology 2012, 11, 66–81, doi:10.1016/S1474-4422(11)70265-4.

9. Hankey, G.J. B Vitamins for Stroke Prevention. Stroke and Vascular Neurology 2018, 3, 51–58, doi:10.1136/svn-2018-000156.

10. Jadavji, N.M.; Emmerson, J.; Willmore, W.G.; MacFarlane, A.J.; Smith, P. B-Vitamin and Choline Supplementation Increases Neuroplasticity and Recovery after Stroke. Neurobiology of Disease 2017, 103, 89–100.

11. Jadavji, N.; Emmerson, J.T.; Shanmugalingam, U.; Willmore, W.G.; Macfarlane, A.J.; Smith, P.D. A Genetic Deficiency in Folic Acid Metabolism Impairs Recovery after Ischemic Stroke. Experimental neurology 2018, 309, 14–22.

12. Abato, J.E.; Moftah, M.; Cron, G.O.; Smith, P.D.; Jadavji, N.M. Methylenetetrahydrofolate Reductase Deficiency Alters Cellular Response after Ischemic Stroke in Male Mice. Nutritional Neuroscience 2020, doi:10.1080/1028415X.2020.1769412.

13. Poole, J.; Jasbi, P.; Pascual, A.S.; North, S.; Kwatra, N.; Weissig, V.; Gu, H.; Bottiglieri, T.; Jadavji, N.M. Reduced Stroke Outcome in Old-Aged Female Mice Maintained on a Dietary Vitamin B12 Deficiency 2022, 2022.04.04.487028.

14. Clementson, M.; Hurley, L.; Coonrod, S.; Bennett, C.; Marella, P.; Pascual, A.S.; Pull, K.; Wasek, B.; Bottiglieri, T.; Malysheva, O.; et al. Maternal Dietary Deficiencies in Folic Acid or Choline Worsen Stroke Outcomes in Adult Male and Female Mouse Offspring. Neural Regen Res 2023, 18, 2443–2448, doi:10.4103/1673-5374.371375.

15. Hurley, L.; Jauhal, J.; Ille, S.; Pull, K.; Malysheva, O.V.; Jadavji, N.M. Maternal Dietary Deficiencies in Folic Acid and Choline Result in Larger Damage Volume, Reduced Neuro-Degeneration and -Inflammation and Changes in Choline Metabolites after Ischemic Stroke in Middle-Aged Offspring. Nutrients 2023, 15, 1556, doi:10.3390/nu15071556.

16. Pull, K.; Folk, R.; Kang, J.; Jackson, S.; Gusek, B.; Esfandiarei, M.; Jadavji, N.M. Impact of Maternal Dietary Folic Acid or Choline Dietary Deficiencies on Vascular Function in Young and Middle-Aged Female Mouse Offspring after Ischemic Stroke. Am J Physiol Heart Circ Physiol 2023, 325, H1354–H1359, doi:10.1152/ajpheart.00502.2023.

17. Mbs, G.B.Y.; Wasek, B.; Bottiglieri, T.; Malysheva, O.; Caudill, M.A.; Jadavji, N.M. Dietary Vitamin B12 Deficiency Impairs Motor Function and Changes Neuronal Survival and Choline Metabolism after Ischemic Stroke in Middle-Aged Male and Female Mice. Nutr Neurosci 2023, 1–10, doi:10.1080/1028415X.2023.2188639.

18. Imbard, A.; Benoist, J.-F.; Blom, H.J. Neural Tube Defects, Folic Acid and Methylation. Int J Environ Res Public Health 2013, 10, 4352–4389, doi:10.3390/ijerph10094352.

19. Virdi, S.; Jadavji, N.M. The Impact of Maternal Folates on Brain Development and Function after Birth. Metabolites 2022, 12, 876, doi:10.3390/metabo12090876.

20. Zeisel, S.H. Choline: Critical Role During Fetal Development and Dietary Requirements in Adults. Annual Review of Nutrition 2006, 26, 229–250, doi:10.1146/annurev.nutr.26.061505.111156.

21. Clare, C.E.; Brassington, A.H.; Kwong, W.Y.; Sinclair, K.D. One-Carbon Metabolism: Linking Nutritional Biochemistry to Epigenetic Programming of Long-Term Development. Annual Review of Animal Biosciences 2019, 7, annurev-animal-020518-115206, doi:10.1146/annurev-animal-020518-115206.

22. Barker, D.J.P.; Godfrey, K.M.; Gluckman, P.D.; Harding, J.E.; Owens, J.A.; Robinson, J.S. Fetal Nutrition and Cardiovascular Disease in Adult Life. The Lancet 1993, 341, 938–941, doi:10.1016/0140-6736(93)91224-A.

23. Barker, D. Intrauterine Programming of Coronary Heart Disease and Stroke. Acta Paediatrica 1997, 86, 178–182, doi:10.1111/j.1651-2227.1997.tb18408.x.

24. Pitkin, R.M. Folate and Neural Tube Defects. Am J Clin Nutr 2007, 85, 285S–288S, doi:10.1093/ajcn/85.1.285S.

25. Shaw, G.M.; Carmichael, S.L.; Yang, W.; Selvin, S.; Schaffer, D.M. Periconceptional Dietary Intake of Choline and Betaine and Neural Tube Defects in Offspring. Am J Epidemiol 2004, 160, 102–109, doi:10.1093/aje/kwh187.

26. Nyaradi, A.; Li, J.; Hickling, S.; Foster, J.; Oddy, W.H. The Role of Nutrition in Children’s Neurocognitive Development, from Pregnancy through Childhood. Front Hum Neurosci 2013, 7, 97, doi:10.3389/fnhum.2013.00097.

27. Georgieff, M.K. Nutrition and the Developing Brain: Nutrient Priorities and Measurement. Am J Clin Nutr 2007, 85, 614S–620S, doi:10.1093/ajcn/85.2.614S.

28. Georgieff, M.K.; Ramel, S.E.; Cusick, S.E. Nutritional Influences on Brain Development. Acta Paediatr 2018, 107, 1310–1321, doi:10.1111/apa.14287.

29. Caffrey, A.; McNulty, H.; Rollins, M.; Prasad, G.; Gaur, P.; Talcott, J.B.; Witton, C.; Cassidy, T.; Marshall, B.; Dornan, J.; et al. Effects of Maternal Folic Acid Supplementation during the Second and Third Trimesters of Pregnancy on Neurocognitive Development in the Child: An 11-Year Follow-up from a Randomised Controlled Trial. BMC Med 2021, 19, 73, doi:10.1186/s12916-021-01914-9.

30. Thornburg, K.L.; O’Tierney, P.F.; Louey, S. Review: The Placenta Is a Programming Agent for Cardiovascular Disease. Placenta 2010, 31 Suppl, S54–59, doi:10.1016/j.placenta.2010.01.002.

31. Lee, H.S.; Kim, W.J. The Role of Matrix Metalloproteinase in Inflammation with a Focus on Infectious Diseases. Int J Mol Sci 2022, 23, 10546, doi:10.3390/ijms231810546.

32. Lavrova, A.I.; Esmedljaeva, D.S.; Belik, V.; Postnikov, E.B. Matrix Metalloproteinases as Markers of Acute Inflammation Process in the Pulmonary Tuberculosis. Data 2019, 4, 137, doi:10.3390/data4040137.

33. Candelario-Jalil, E.; Yang, Y.; Rosenberg, G.A. Diverse Roles of Matrix Metalloproteinases and Tissue Inhibitors of Metalloproteinases in Neuroinflammation and Cerebral Ischemia. Neuroscience 2009, 158, 983–994, doi:10.1016/j.neuroscience.2008.06.025.

34. Tejima, E.; Guo, S.; Murata, Y.; Arai, K.; Lok, J.; van Leyen, K.; Rosell, A.; Wang, X.; Lo, E.H. Neuro-protective Effects of Overexpressing Tissue Inhibitor of Metalloproteinase TIMP-1. Journal of neurotrauma 2009, 26, 1935–1941, doi:10.1089/neu.2009.0959.

35. Fujimoto, M.; Takagi, Y.; Aoki, T.; Hayase, M.; Marumo, T.; Gomi, M.; Nishimura, M.; Kataoka, H.; Hashimoto, N.; Nozaki, K. Tissue Inhibitor of Metalloproteinases Protect Blood-Brain Barrier Disruption in Focal Cerebral Ischemia. J Cereb Blood Flow Metab 2008, 28, 1674–1685, doi:10.1038/jcbfm.2008.59.

36. Reeves, P.G.; Nielsen, F.H.; Fahey, G.C. AIN-93 Purified Diets for Laboratory Rodents: Final Report of the American Institute of Nutrition Ad Hoc Writing Committee on the Reformulation of the AIN-76A Rodent Diet. The Journal of nutrition 1993, 123, 1939–1951.

37. Jadavji, N.M.; Deng, L.; Malysheva, O.; Caudill, M.A.; Rozen, R. MTHFR Deficiency or Reduced Intake of Folate or Choline in Pregnant Mice Results in Impaired Short-Term Memory and Increased Apoptosis in the Hippocampus of Wild-Type Offspring. Neuroscience 2015, 300, 1–9, doi:10.1016/j.neuroscience.2015.04.067.

38. Jadavji, N.M.; Farr, T.; Khalil, A.; Boehm-Sturm, P.; Foddis, M.; Harms, C.; Füchtemeier, M.; Dirnagl, U. Elevated Levels of Plasma Homocysteine, Deficiencies in Dietary Folic Acid and Uracil-DNA Glycosylase Impair Learning in a Mouse Model of Vascular Cognitive Impairment. Behavioural brain research 2015, 283, 215–226, doi:10.1016/j.bbr.2015.01.040.

39. Clementson, M.; Hurley, L.; Coonrod, S.; Bennett, C.; Marella, P.; Pascual, A.S.; Pull, K.; Wasek, B.; Bottiglieri, T.; Malysheva, O.; et al. Maternal Dietary Deficiencies in Folates or Choline during Pregnancy and Lactation Worsen Stroke Outcome in 3-Month-Old Male and Female Mouse Offspring 2022, 2022.09.28.509960.

40. Jadavji, N.M.; Mosnier, H.; Kelly, E.; Lawrence, K.; Cruickshank, S.; Stacey, S.; McCall, A.; Dhatt, S.; Arning, E.; Bottiglieri, T.; et al. One-Carbon Metabolism Supplementation Improves Outcome after Stroke in Aged Male MTHFR-Deficient Mice. Neurobiology of Disease 2019, 132, doi:10.1016/j.nbd.2019.104613.

41. Yahn, G.; Abato, J.; Jadavji, N. Role of Vitamin B12 Deficiency in Ischemic Stroke Risk and Outcome. Neural Regeneration Research 2021, 16, doi:10.4103/1673-5374.291381.

42. Jadavji, N.M.; Emmerson, J.T.; Shanmugalingam, U.; MacFarlane, A.J.; Willmore, W.G.; Smith, P.D. A Genetic Deficiency in Folic Acid Metabolism Impairs Recovery after Ischemic Stroke. Exp Neurol 2018, 309, 14–22, doi:10.1016/j.expneurol.2018.07.014.

43. Jadavji, N.M.; Emmerson, J.T.; MacFarlane, A.J.; Willmore, W.G.; Smith, P.D. B-Vitamin and Choline Supplementation Increases Neuroplasticity and Recovery after Stroke. Neurobiol Dis 2017, 103, 89–100, doi:10.1016/j.nbd.2017.04.001.

44. Jadavji, N.M.; Mosnier, H.; Kelly, E.; Lawrence, K.; Cruickshank, S.; Stacey, S.; McCall, A.; Dhatt, S.; Arning, E.; Bottiglieri, T.; et al. One-Carbon Metabolism Supplementation Improves Outcome after Stroke in Aged Male MTHFR-Deficient Mice. Neurobiol Dis 2019, 132, 104613, doi:10.1016/j.nbd.2019.104613.

45. Löffek, S.; Schilling, O.; Franzke, C.-W. Series “Matrix Metalloproteinases in Lung Health and Disease”: Biological Role of Matrix Metalloproteinases: A Critical Balance. Eur Respir J 2011, 38, 191–208, doi:10.1183/09031936.00146510.

46. Roth, W.; Mohamadzadeh, M. Vitamin B12 and Gut-Brain Homeostasis in the Pathophysiology of Ischemic Stroke. EBioMedicine 2021, 73, 103676, doi:10.1016/j.ebiom.2021.103676.

47. Mbs, G.B.Y.; Wasek, B.; Bottiglieri, T.; Malysheva, O.; Caudill, M.A.; Jadavji, N.M. Dietary Vitamin B12 Deficiency Impairs Motor Function and Changes Neuronal Survival and Choline Metabolism after Ischemic Stroke in Middle-Aged Male and Female Mice. Nutritional Neuroscience 2024, 27, 300–309, doi:10.1080/1028415X.2023.2188639.

48. Aslam, A.; Misbah, S.A.; Talbot, K.; Chapel, H. Vitamin E Deficiency Induced Neurological Disease in Common Variable Immunodeficiency: Two Cases and a Review of the Literature of Vitamin E Deficiency. Clinical Immunology 2004, 112, 24–29, doi:10.1016/j.clim.2004.02.001.

49. Jeng, K.; Yang, C.; Siu, W.; Tsai, Y.; Liao, W.; Kuo, J. Supplementation with Vitamins C and E Enhances Cytokine Production by Peripheral Blood Mononuclear Cells in Healthy Adults. The American Journal of Clinical Nutrition 1996, 64, 960–965, doi:10.1093/ajcn/64.6.960.

50. Green, J.A.; Dholakia, S.; Janczar, K.; Ong, C.W.; Moores, R.; Fry, J.; Elkington, P.T.; Roncaroli, F.; Friedland, J.S. Mycobacterium Tuberculosis-Infected Human Monocytes down-Regulate Microglial MMP-2 Secretion in CNS Tuberculosis via TNFα, NFκB, P38 and Caspase 8 Dependent Pathways. J Neuroinflammation 2011, 8, 46, doi:10.1186/1742-2094-8-46.

51. Owolabi, U.S.; Amraotkar, A.R.; Coulter, A.R.; Singam, N.S.V.; Aladili, B.N.; Singh, A.; Trainor, P.J.; Mitra, R.; DeFilippis, A.P. Change in Matrix Metalloproteinase 2, 3, and 9 Levels at the Time of and after Acute Atherothrombotic Myocardial Infarction. J Thromb Thrombolysis 2020, 49, 235–244, doi:10.1007/s11239-019-02004-7.

