## Supplementary Materials: Table S1: Mean and Standard Error of Mean data. for "Maternal dietary deficiency in choline reduced levels of MMP-2 levels in blood and brain tissue of male offspring mice"

Plasma

| | | | | |
|---|---|---|---|---|
| Figure 2A | MMP-2 | Female | Mean | SEM |
| | | CD | 322.761 | 20.32 |
| | | FADD | 365.233 | 22.72 |
| | | ChDD | 331.491 | 16.305 |
| | | Male | | |
| | | CD | 317.179 | 14.668 |
| | | FADD | 358.908 | 19.894 |
| | | ChDD | 277.636 | 12.366 |
| Figure 2B | MMP-9 | Female | Mean | SEM |
| | | CD | 27.121 | 3.87 |
| | | FADD | 32.011 | 4.278 |
| | | ChDD | 29.876 | 4.476 |
| | | Male | | |
| | | CD | 29.511 | 3.683 |
| | | FADD | 26.323 | 2.639 |
| | | ChDD | 24.448 | 2.586 |

Brain Tissue

| | | | | |
|---|---|---|---|---|
| Figure 3 | MMP-2 | Female | Mean | SEM |
| | | CD | 285.625 | 39.76 |
| | | FADD | 208.611 | 25.322 |
| | | ChDD | 236.083 | 38.656 |
| | | Male | | |
| | | CD | 317.743 | 24.185 |
| | | FADD | 231.333 | 41.939 |
| | | ChDD | 175.444 | 27.215 |
| Figure 4 | MMP-2 | Female | Mean | SEM |
| | | CD | 340.778 | 30.642 |
| | | FADD | 222 | 35.533 |
| | | ChDD | 197.022 | 88.994 |
| | | Male | | |
| | | CD | 247.444 | 72.204 |
| | | FADD | 207.667 | 27.713 |
| | | ChDD | 238.667 | 18.667 |
